# Competitive Tolerance And Yield Response Of Maize, Cowpea And Tomato In Mixed Culture And Pure Stand

**DOI:** 10.1101/344747

**Authors:** Ezekiel Dare Olowolaju, Gideon Olarewaju Okunlola, Adekunle AJayi Adelusi

**Author notes:** Corresponding author’s. **Addresses of the institutions where the work was carried out**: Obafemi Awolowo University, Faculty of Science, Department of Botany, Ile-Ife, Nigeria.

## Abstract

This study investigated the competitive effect and yield response of maize, cowpea and tomato in mixed and pure stand.

The experiment was carried out under a screenhouse to minimize extraneous factors such as pests and rodents using a randomized complete block design (RCBD). The treatments where T1 = sole maize (SM); T2 = Maize intercrop with cowpea (MC); T3 = Maize intercrop with tomato (MT); T4 = Sole tomato (ST); T5 = Tomato intercrop with Cowpea (TC); T6 = Sole cowpea (SC); T7 = Maize intercrop with cowpea and tomato (MCT). At harvest the yields parameters of maize, tomato and cowpea were taken. Some of the indices for intensity of competition were computed using standard formulas and statistical analysis was performed using statistical analytical software SAS version 9.12.

The result showed that the yield of maize, cowpea and tomato were enhanced in the intercropped than the sole crops. Most of the intensity of competition indices of maize, tomato and cowpea in the mixed cultured were greater than 1, while that of the sole culture equals 1.

The study concluded that competition for shared resources in the mixed culture of tomato, maize and cowpea enhanced their yield.

## Introduction

Competitive response or tolerance is the ability of an organism to continue to perform relatively well in the presence of competitors (Goldberg and Werner, 1983; Miller and Werner, 1987). The response to competition entails the ability of target plants to avoid being suppressed by their companions (Goldberg and Werner, 1983). When crops are subjected to strong competition stress, the physiological characteristics of growth, yield and development are usually changed. This was as a result of differences in the response of the competing plants in the use of environmental resources such as light, water, space and nutrients (Olowolaju and Adelusi, 2017). Competitive stress created by competition in plant stands may be expressed by increased mortality, reduced seed production, and reduced growth rate (Board, 2000).

Yield in the multiple cropping occurs when components crops differ in their use of growth resources in such a way that when they are grown in combination they are better able to complement each other and make better use of resources than when grown in separately. The component crops are not competing for exactly the same resources (space and time) because component crops differ in their nutrients requirements which can readily exploits and their ability to extract them from the soil. Where the total quantities of resource captured is relatively similar, the efficiency of utilization of the resources captured is increased in the intercrops compared to the sole crops. The advantages of intercropping to competition of plants of the same or different species is the production of greater yield on a given space (Olowolaju and Okunlola, 2017) by making more efficient use of the available growth resources using a mixture of crops of different rooting ability, canopy structure, height, and nutrient requirements based on the complementary utilization of growth resources by the component crops in a limited space (Lammerts van Bueren *et al.,* 2002).

Yield can be considered as a function of four basic factors which include seed mass, number of seeds pod, number of pods plant, and number of plants per given area. Identifying which yield components contribute the most to yield and yield compensation under given crop management situations would help to understand necessary management to achieve optimal yields. Moreover, an increased understanding of how yield components and growth dynamic factors regulate maize, cowpea and tomato yield in response to plant population could improve cultivar development to optimize yield, that could also provide producers with indicators of optimal intercropping system that will improve the yield components’ of maize, cowpea and tomato. The yield advantage of intercropping of more than two crops has not been so marked. Therefore, the objective of this study is to investigate the yield response of interplant competition of maize, cowpea and tomato in pure and mixed stand.

## MATERIALS AND METHODS

The Seeds of cowpea (IT07K-38-33), maize (2008 DTMA-YSTR) and tomato (ROMA VF) were utilized in this experiment. These seeds were collected from the Department of Crop Production and Protection, Faculty of Agriculture, Obafemi Awolowo University, Ile Ife, Osun state. A screenhouse was constructed to minimize extraneous factors such as pests and rodents, supply of water other than the amount specifically applied. The mean daily temperature under the screenhouse was taken with the aid of a thermometer. The intensity of light was also determined using a digital luxmeter LX 1000. Relative humidity was measured using a hygrometer.

### Raising of Seedlings

Forty two bowls were obtained (of 38cm in diameter and 5.5 cm in height). Holes of about 3mm each were bored at the bottom of the bowls. This is to allow for proper drainage and prevent water logging during the course of the experiment. The bowls were filled near brim with 10 kg of the analyzed soil. The seeds of cowpea, maize and tomato were then planted at a depth of about 3 mm below the soil. The seeds were sown at the rate of six seeds per pot in the monoculture, while in the pots designed for the mixed culture of maize and cowpea, maize and tomato, cowpea and tomato, three seeds of each plant were sown. Two seeds of each plant were sown in the pots with the three crops. The bowls were then supplied with 500 ml of tap water in the morning and 500 ml of tap water in the evening until the seedlings become fully established.

### Experimental layout

The experiment was laid out in a Randomized Complete Block Design (RCBD) with six replicates The seedlings were divided into seven regimes which include the following: T1 = sole maize (SM); T2 = Maize intercrop with cowpea (MC); T3 = Maize intercrop with tomato (MT); T4 = Sole tomato (ST); T5 = Tomato intercrop with cowpea (TC); T6 = Sole cowpea (SC); T7 = Maize intercrop with cowpea and tomato (MCT). All the groups of plants were made to receive 500 ml of water every morning and evening throughout the experimental period.

### Determination of Yield Parameters

At harvest, the following yield parameters were taken:

### Number of pod per plant

Pods from randomly selected ten plants from each regime of cowpea was counted and the average was taken as the number of pods per plant.

### Number of seeds in pod per plant

The seeds from ten representative mature pods were separated and counted and the average was taken as the number of pods per plant.

### Pod length

The length of pod of cowpea from randomly selected ten plants from each regimes was measured with the aid of calibrated metric rule and averages were taken as the number of pods per plant.

### Number of cob per plant

Cobs from randomly selected ten plants from each regime of maize were counted and the averages were taken as the number of cob per plant.

### Number of seeds in cob

The seeds from randomly selected matured cobs of maize from each regime were shelled and counted and the averages were taken as the number of seeds in cob per plant.

### Cob length

The length of cobs of maize from randomly selected ten plants from each regime was counted and averages were taken.

### Number of fruits per plant

The fruits from randomly selected mature fruits of tomato from each regime were counted and the averages were taken.

### Seed yield/Kg per pot

Seeds were separated by shelling from cobs and pods, were oven dried at 35 degree Celsius for three days, weighed and the seed weight were recorded and computed to kg/pot. This was used to determine the crop yield of maize and cowpea.

### Fruit yield/kg per pot

The fruits were harvested from randomly selected plants. The diameter was measured with vernier caliper and fruit weight was recorded and computed to kg/pot. This was used to determine the crop yield of tomato.

### Indices of the Intensity of Competition

These were determined by relative reproductive rate (RRR), relative yield of mixture (RYM), relative competition intensity (RCI) and relative resource total (RRT).

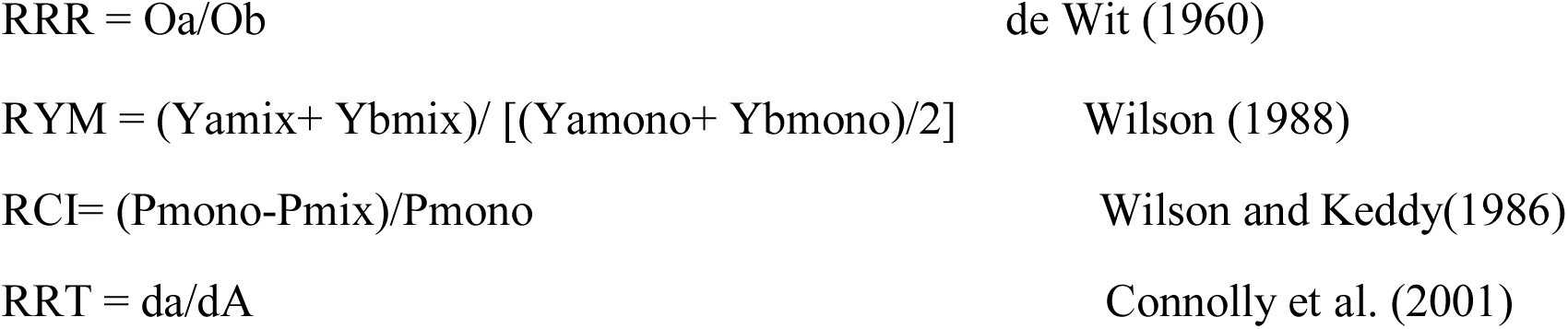

Where Y is performance per unit area (with Y = P × d; (where P is performance per plant (yield), d is planting density); O is number of seeds produced per plant, a and b for plant in pure and mixed stand; Pmono and Pmix is relative growth rate of plant in the pure and mixed stand. dA is the density of plant in the pure stand, da is the density of plant in the mixed stand

### Statistical Analysis

Statistical analysis was performed using statistical analytical software SAS version 9.0. A one way analysis of variance (ANOVA) was carried out to investigate the effect of competition stress on the growth and yield parameters. Post hoc setting testing was carried out using Bonferroni to separate the significance means at 0.05 confidence limit (alpha level) for the mean.

## RESULTS AN DISCUSSION

### The Yield of Maize in Sole Culture and in the Mixed Culture

The cob length and cob weight (Table 4) of T3-plants was statistically higher and that of T7-plants was statistically lower (P>0.05) to other treatments. There was no significant difference (P>0.05) in the cob length and cob weight of all the treatments. The higher cob length and cob weight observed in T3-plants might be attributed to allocation of resource use in this treatments compare to other treatments that mature and flower earlier

The number of seeds per cob (Table 4) was statistically higher in T7-plantsand was statistically lower (P>0.05) in T2-plants compare to other treatments. The number of seeds per cob of T2-plants was statistically different (P>0.05) from the number of seeds per cob of T2-plants. The number of seeds per cob of T2-plants and T7-plants was not statistically different (P>0.05) from the number of seeds per cob of T1-plants and T3-plants (P>0.05). The number of seeds per cob which was higher in T7-plants compared to other treatments might be due to relative neighbor effects which promote early maturity and thus minimum time was available for seed setting and development due to competition for shared resource among the components

The seed weights of maize plant (Table 4) was statistically higher in T7-palnts and was statistically lower (P>0.05) in T2-plants compare to other treatments. The seed weight of T2-plants was statistically different from the weight of T2-plants. There was no statistical different in the seed weights of all the treatments (P>0.05). The highest seed weight recorded in T7-plants might be attributed to the acquisition of more resource use in the mixed culture.

### The Yield of Cowpea in Sole Culture and in the Mixed Culture

The pod length of T7-plants (Table 6) was statistically higher (P>0.05) and the pod length of T6-plants was statistically lower than the other treatments. The pod weight of T7-plants was statistically higher (P>0.05) and the pod weight of T2-plants was statistically lower to other treatments. There was no significant difference (P>0.05) in the pod length and pod weight of all the treatments. The pod length and pod weight of T7-plants which was statistically higher may be attributed to the ability of cowpea to use the available resources when competing with other crops; meanwhile, cowpea was the dominant crop in the intercrop.

The number of seed per pod of T6-plants (Fig. 12) was statistically higher (P>0.05) in all the treatments from the 63^rd^ day after emergence to the end of the experimental period. There was significant difference (P>0.05) in the number of seed per pod of all the treatments from the day of emergence to the end of the experiment. The higher number of seeds per pod observed in T6- is in agreement with the study of Abraha (2013), who observed higher number of seeds per pod in the monoculture of cowpea compared to the intercropped.

The seed weight of T5-plants (Table 6) was statistically higher (P>0.05) and the seed weight of T6-plants was statistically lower than the other treatments. There was significant difference (P>0.05) in the seed weight of all the treatments. The higher seed weight observed in T5-plants and lower seed weight in T7-plants may be due to availability of photosynthate for seed development in T5-plants

### The Yield of Tomato in Sole Culture and in the Mixed Culture

The number of fruits of T4-plants (Fig. 11.2) was statistically higher and the number of fruits of T7-plants was statistically lower (P>0.05) to other treatments from the day of emergence to the end of the experimental period. There was no significant difference in the number of fruits of all the treatments from the day of emergence to the end of the experimental period (P>0.05). The number of fruits of T4-plants which was statistically higher than the other treatments reflect an interspecific relationship of mutual inhibition in which crops in their mixtures at the various planting patterns yielded less than their potential (expected) yields in monoculture (Abraha, 2003).

The fruit length (diameter) of T7-plants (Table 5) was the highest and the fruit length of T3-plants was the lowest at P>0.05. The fruit length of all the treatments was statistically different from one another (P>0.05). T3-plants which recorded the lowest was attributed to the height-to-crown and size asymmetric of maize plants to tomato plants (Schmitt and Wulf, 1993).This might also be attributed to shadowing and suppression nature of the cereal mixture over the component crop.

The fruit weight of T5-plants (Table 5) was statistically higher (P>0.05) and the fruit weight of T3-plants was statistically lower (P>0.05) to other treatments. The result of the ANOVA shows significant difference (P>0.05) in the fruit weight of all the treatments. T5-plants which had the highest fruit weight indicated that T5-plants shows a possibility to capture available resources in the mixture.

### Indices of the Intensity of Competition of Maize in Sole Culture and in the Mixed Culture

The result from the table 2 shows the RRR of T2-plants and T3-plants were greater than 1, RRR of T1-plants equals 1 and RRR of T1-plants was less than 1. The RRR of T1-plants was significantly different from the other treatments at P≤0.05.

**Table 1:**
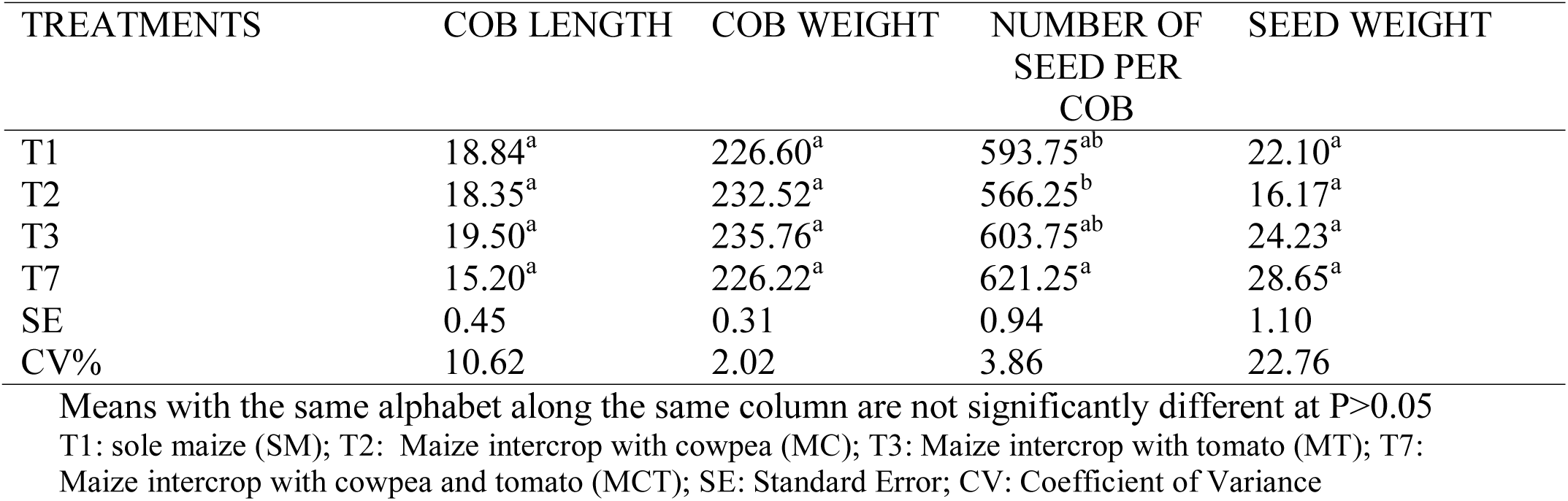
Cob length, cob weight, number of seed per cob and seed weight of maize in sole culture and mixed culture

| TREATMENTS | COB LENGTH | COB WEIGHT | NUMBER OF SEED PER COB | SEED WEIGHT |
| --- | --- | --- | --- | --- |
| T1 | 18.84<sup>a</sup> | 226.60<sup>a</sup> | 593.75<sup>ab</sup> | 22.10<sup>a</sup> |
| T2 | 18.35<sup>a</sup> | 232.52<sup>a</sup> | 566.25<sup>b</sup> | 16.17<sup>a</sup> |
| T3 | 19.50<sup>a</sup> | 235.76<sup>a</sup> | 603.75<sup>ab</sup> | 24.23<sup>a</sup> |
| T7 | 15.20<sup>a</sup> | 226.22<sup>a</sup> | 621.25<sup>a</sup> | 28.65<sup>a</sup> |
| SE | 0.45 | 0.31 | 0.94 | 1.10 |
| CV% | 10.62 | 2.02 | 3.86 | 22.76 |

Means with the same alphabet along the same column are not significantly different at $P>0.05$
T1: sole maize (SM); T2: Maize intercrop with cowpea (MC); T3: Maize intercrop with tomato (MT); T7: Maize intercrop with cowpea and tomato (MCT); SE: Standard Error; CV: Coefficient of Variance

**Table 2:** Pod length, pod weight, number of seed per pod and seed weight of cowpea in sole and mixed culture

| TREATMENTS | POD LENGTH | POD WEIGHT | NUMBER OF SEED PERPOD | SEED WEIGHT |
| --- | --- | --- | --- | --- |
| T2 | 14.18<sup>a</sup> | 3.08<sup>a</sup> | 11.75<sup>a</sup> | 11.38<sup>b</sup> |
| T5 | 16.18<sup>a</sup> | 3.48<sup>a</sup> | 13.00<sup>a</sup> | 12.67<sup>a</sup> |
| T6 | 16.00<sup>a</sup> | 3.61<sup>a</sup> | 13.25<sup>a</sup> | 8.72<sup>d</sup> |
| T7 | 16.58<sup>a</sup> | 3.90<sup>a</sup> | 11.50<sup>a</sup> | 10.68<sup>c</sup> |
| SE | 0.27 | 0.09 | 0.25 | 0.50 |
| CV% | 6.77 | 9.66 | 7.094 | 15.18 |

Means with the same alphabet along the same column are not significantly different at $P>0.05$ T2: Maize intercrop with cowpea (MC); T5: Tomato intercrop with cowpea (TC); T6: Sole cowpea (SC); T7: Maize intercrop with cowpea and tomato (MCT); SE: Standard Error; CV: Coefficient of Variance

**Table 3:** Fruit length and fruit weight of tomato in sole culture and mixed culture

| TREATMENTS | FRUIT LENGTH | FRUIT WEIGHT |
| --- | --- | --- |
| T3 | 2.50<sup>c</sup> | 14.48<sup>d</sup> |
| T4 | 3.20<sup>b</sup> | 20.43<sup>b</sup> |
| T5 | 4.50<sup>a</sup> | 28.74<sup>a</sup> |
| T7 | 4.00<sup>a</sup> | 17.32<sup>c</sup> |
| SE | 0.47 | 1.37 |
| CV% | 24.82 | 30.45 |

Means with the same alphabet along the same column are not significantly different at $P>0.05$ ; T3: Maize intercrop with tomato (MT); T4: Sole tomato (ST); T5: Tomato intercrop with cowpea (TC); T7: Maize intercrop with cowpea and tomato (MCT); SE: Standard Error; CV: Coefficient of Variance

**Table 4:**
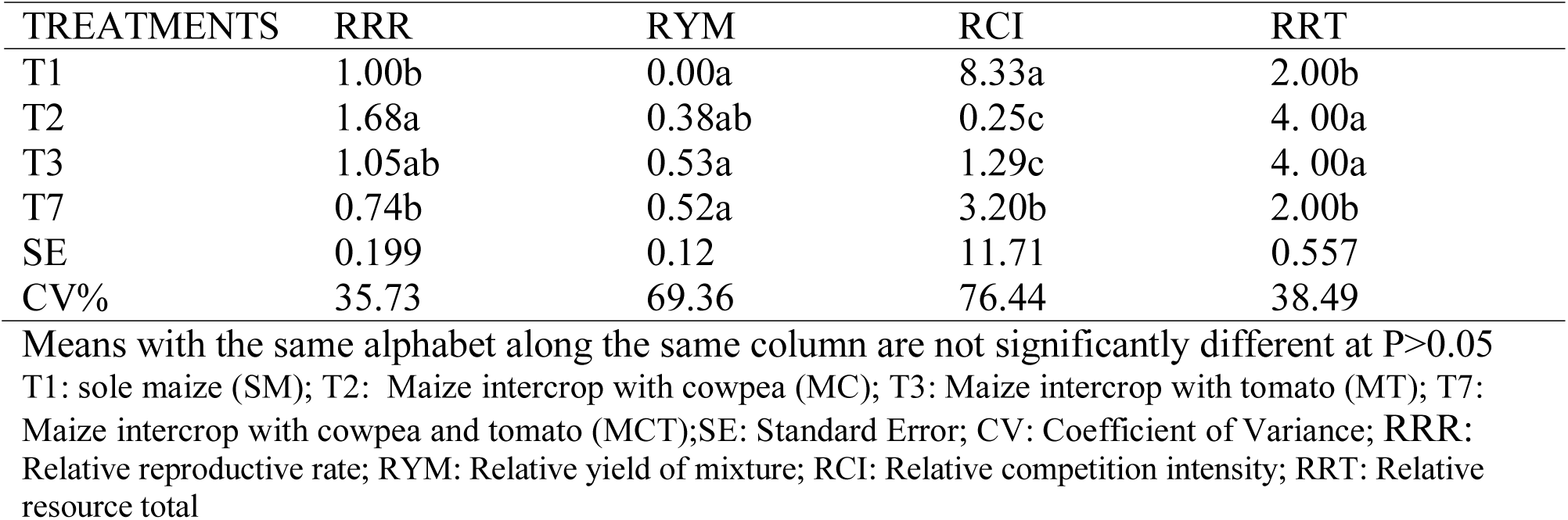
Indices of the Intensity of Competition of Maize in Sole Culture and in the Mixed Culture

**Table 5:** Indices of the Intensity of Competition of Tomato in Sole Culture and in the Mixed Culture

| TREATMENTS | RRR | RYM | RCI | RRT |
| --- | --- | --- | --- | --- |
| T3 | 1.23a | 0.50a | 0.85a | 1.64a |
| T4 | 1.00a | 0.00a | 1.08a | 2.00a |
| T5 | 3.20a | 0.50a | 1.17a | 2.43a |
| T7 | 0.00a | 0.50a | 0.12a | 1.38a |
| SE | 0.13 | 0.93 | 0.24 | 0.93 |
| CV% | 24.47 | 100 | 59.14 | 100 |

Means with the same alphabet along the same column are not significantly different at $P > 0.05$ ; T3: Maize intercrop with tomato (MT); T4: Sole tomato (ST); T5: Tomato intercrop with cowpea (TC); T7: Maize intercrop with cowpea and tomato (MCT); SE: Standard Error; CV: Coefficient of Variance; RRR: Relative reproductive rate; RYM: Relative yield of mixture; RCI: Relative competition intensity; RRT: Relative resource total

**Table 6:** Indices of the Intensity of Competition Cowpea in Sole Culture and in the Mixed Culture

| TREATMENTS | RRR | RYM | RCI | RRT |
| --- | --- | --- | --- | --- |
| T2 | 1.13a | 0.11a | 0.78c | 4.00a |
| T5 | 1.065a | 0.03a | 0.94b | 4.00a |
| T6 | 1.00a | 0.00a | 2.20a | 2.00b |
| T7 | 1.22a | 0.15a | 1.57ab | 2.00b |
| SE | 0.047 | 0.0034 | 0.32 | 0.577 |
| CV% | 8.5 | 95.81 | 47.25 | 38.49 |

Means with the same alphabet along the same column are not significantly different at $P>0.05$ T2: Maize intercrop with cowpea (MC); T5: Tomato intercrop with cowpea (TC); T6: Sole cowpea (SC); T7: Maize intercrop with cowpea and tomato (MCT); SE: Standard Error; CV: Coefficient of Variance; RRR: Relative reproductive rate; RYM: Relative yield of mixture; RCI: Relative competition intensity; RRT: Relative resource total

The RYM of all the treatments were less than 1. There was no significant difference in the RYM of all the treatments at P≤0.05.

The RRT of all the treatments were greater than 1 and there was no significant difference in the RRT of all the treatments at P≤0.05.

The RCI of T1-, T3- and T7-plants were greater than 1, while that of T2-plants was less than 1. The RCI of T-plants and T7-plants were significantly different from each other and from T2-plants and T3-plants at P<0.05.

### Indices of the Intensity of Competition of Tomato in Sole Culture and in the Mixed Culture

The result from the table 2 shows the RRR of T3-plants and T5-plants were greater than 1, RRR of T4-plants equals 1 and RRR of T7-plants was less than 1. There was no significant difference in the RRR OF all the treatments at P≤0.05.

The RYM of all the treatments were less than 1. There was no significant difference in the RYM of all the treatments at P≤0.05.

The RRT of all the treatments were greater than 1 and there was no significant difference in the RRT of all the treatments at P≤0.05

The RCI of T4- and T5-plants were greater than 1, while that of T3-plants and T7-plants were less than 1. There was no significant difference in the RRT of all the treatments at P≤0.05

### Indices of the Intensity of Competition of Cowpea in Sole Culture and in the Mixed Culture

The result from the table 2 shows the RRR of T2-, T5-plants and T7-plants were greater than 1, while RRR of T6-plants equals 1. There was no significant difference in the RRR of all the treatments at P≤0.05.

The RYM of all the treatments were less than 1. There was no significant difference in the RYM of all the treatments at P≤0.05.

The RRT of all the treatments were greater than 1 and there was no significant difference in the RRT of all the treatments of T2- and T5-plants, T6-plants and T7-plants but these groups were significantly different from each other at P≤0.05.

The RCI of T6- and T7-plants were greater than 1, while that of T2-plants and T5-plants were less than 1. There was significant difference in the RRT of all T2-, T5-plants and T6-plants the treatments at P≤0.05

The intensity of competition of maize, tomato and cowpea plants in all the treatments which were greater than 1 in the mixed stand shows that these plants were able to compliments each other indicating yield advantage in the mixed culture. These also show that these plants have competitive effects on each other, thereby enhancing their yield performance. The intensity of competition of these plants in the pure stands which were greater than 1 implies that there was separate interference of maize, tomato and cowpea in the sole culture compared to the mixed culture. The intensity of competition of maize, tomato and cowpea plants in all the treatments which were less than 1 can be inferred that maize plants produced less yield as expected presumably due to inadequate utilization of resources. The intensity of competition of maize plants in all the treatments both in the mixed and pure stand which equals to 1 indicated that competition for shared resources used has no effects on the yield of maize, tomato and cowpea

## CONCLUSION

Due to the reduced yields of maize, cowpea and tomato in sole culture compared to the mixed culture it can be concluded that maize, cowpea and tomato can be grown together for their optimum yield. Depending on the producers objectives for good land management and/or available land, intercropping maize, tomato with cowpea can be practiced to produce more yields.

